# Molecular recognition of M1-linked ubiquitin chains by native and phosphorylated UBAN domains

**DOI:** 10.1101/521815

**Authors:** Lina Herhaus, Henry van den Bedem, Sean Tang, Soichi Wakatsuki, Ivan Dikic, Simin Rahighi

**Author notes:** These authors contributed equally to the manuscript (Lina Herhaus, Henry van den Bedem).

## Abstract

Although the Ub-binding domain in ABIN proteins and NEMO (UBAN) is highly conserved, UBAN-containing proteins exhibit different Ub-binding properties, resulting in their diverse biological roles. Post-translational modifications further control UBAN domain specificity for poly-Ub chains. However, precisely, how the UBAN domain structurally confers such functional diversity remains poorly understood. Here we report crystal structures of ABIN-1 alone and in complex with one or two M1-linked di-Ub chains. ABIN-1 UBAN forms a homo-dimer that provides two symmetrical Ub-binding sites on either side of the coiled-coil structure. Moreover, crystal structures of ABIN1 UBAN in complex with di-Ub chains reveal a concentration-dependency of UBAN/di-Ub binding stoichiometry. Analysis of UBAN/M1-linked di-Ub binding characteristics indicates that phosphorylated S473 in OPTN and its corresponding phospho-mimetic residue in ABIN-1 (E484) are essential for high affinity interactions with M1-linked Ub chains. Also, a phospho-mimetic mutation of A303 in NEMO, corresponding to S473 of OPTN, increases binding affinity for M1-linked Ub chains. These findings are in line with the diverse physiological roles of UBAN domains, as phosphorylation of OPTN UBAN is required to enhance its binding to Ub during mitophagy.

## Introduction

Ubiquitination is a post-translational modification (PTM) in which ubiquitin (Ub) is attached to target proteins through a cascade of enzymatic reactions catalyzed by Ub activating (E1), conjugating (E2) and ligase (E3) enzymes [1, 2]. Target proteins can be mono-ubiquitinated or poly-ubiquitinated by 8 different homotypic chains or mixed and branched chains, in which Ub molecules are attached via any of the M1, K6, K11, K27, K29, K33, K48 or K63 residues [3]. These chains with different linkage types adopt a variety of conformations, and function as signals that are specifically recognized by Ub-binding proteins, which relay the signal to downstream proteins resulting in distinct cellular responses [4].

To date, over 20 different types of Ub-binding domains (UBDs) have been identified [4]. UBAN is a UBD shared by Optineurin (OPTN), NEMO (NF-κB essential modulator) and ABIN1-3 (A20-binding inhibitors of NF-κB) proteins (Fig. 1A) [4-6]. Binding of Ub chains to UBAN domains is an essential step in the regulation of cellular functions of the UBAN-containing proteins. NEMO and ABIN-1 play essential roles in the NF-κB signaling pathway, which regulates gene expression, thereby affecting inflammation, tumorigenesis and immunity and is extensively regulated through M1- and K63-linked ubiquitination [7-11]. NEMO is a component of the inhibitor of κB (IκB) kinase (IKK), that regulates activation of the kinase complex and subsequent activation of the NF-κB signaling pathway. ABIN-1 controls innate immune responses through the NF-κB signaling. It binds to linearly ubiquitinated NEMO and A20 ubiquitin-editing enzyme and promotes deubiquitination and termination of NF-κB signaling [12]. Also, upon tumor necrosis factor (TNF)-α stimulation, ABIN-1 is recruited to the TNF receptor 1 (TNFR1) resulting in the recruitment of A20 and deubiquitination of RIP kinase 1 [13, 14]. OPTN is involved in diverse cellular functions [15]. Autophagic degradation of depolarized mitochondria (mitophagy) and intracellular pathogens (xenophagy) heavily rely on the phosphorylation of OPTN UBAN by the kinase TBK1. Locally accumulated and active TBK1 phosphorylates the UBAN domain of OPTN (on S473) at the phagophore [16] and phosphorylated OPTN (together with other autophagic receptors) bridges damaged mitochondria to LC3-coated phagophores, thereby driving autophagy. Disturbing this pathway through genetic alterations and impairing TBK1 activity or its binding to OPTN have been linked to frontotemporal lobar degeneration (FLD) and amyotrophic lateral sclerosis (ALS) [17-20].

**Figure 1.**
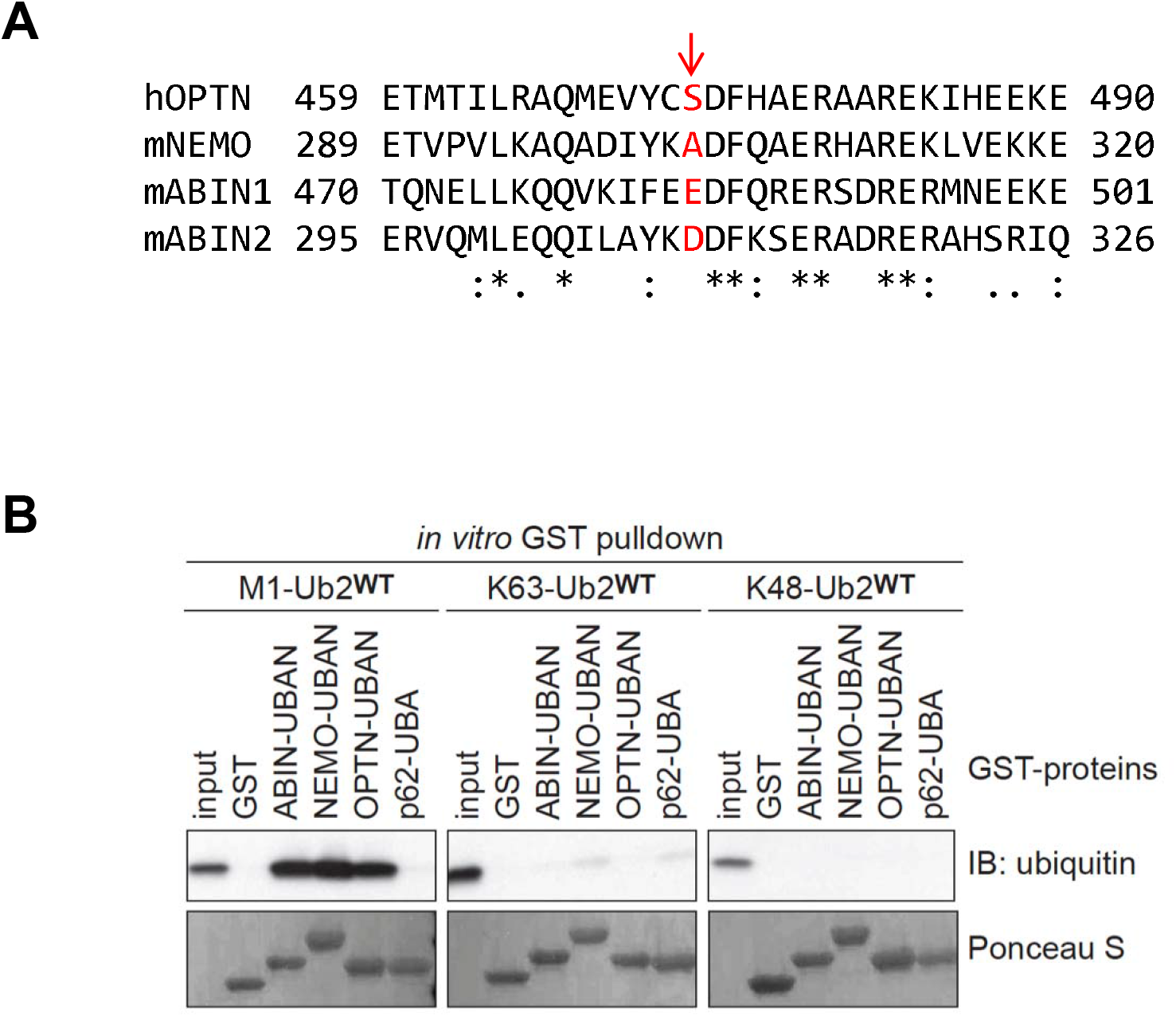
UBAN domains preferentially bind M1-linked di-Ub chains. (**A**) Amino-acid sequence alignment of UBAN domains in hOPTN, mNEMO and mABIN1-3 proteins. The residues modulating di-Ub binding are marked in red. The sequence alignment was performed using Clustal Omega. * indicates positions which have a fully conserved residue. A colon indicates conservation between groups of strongly similar properties (>0.5 Gonnet PAM 250 matrix) and a period indicates conservation between groups of weakly similar properties (0-0.5 Gonnet PAM 250 matrix). (**B**) GST immunoprecipitation of purified UBAN domains of OPTN, ABIN1, NEMO, p62 or GST only with M1-, K63- or K48-linked di-Ub chains.

Crystal structures of various UBAN domains have demonstrated a parallel coiled-coil homo-dimeric structure that provides two binding sites for M1-linked di-Ub chains [21-23]. NEMO UBAN also binds K63-linked di-Ub chains, albeit with about 100-fold lower affinity than M1-linked di-Ub chains [24]. While UBAN domains adopt near identical conformations in binding Ub, they can exhibit different Ub-binding characteristics determined by the cellular role of their host protein. Therefore, understanding the molecular mechanisms that underpin the function of UBAN-containing proteins requires detailed analysis of UBAN/Ub interactions. In this study, we have analyzed the structure of ABIN-1 UBAN and its interactions with M1-linked Ub chains. We have determined crystal structures of ABIN-1 UBAN alone and in complex with M1-linked di-Ub chains and investigated stoichiometry of UBAN/di-Ub chain binding. We have also examined the effects of UBAN phosphorylation on binding of M1-linked Ub chains using molecular dynamics (MD) simulations and *in vitro* binding assays.

## Results

### ABIN-1 UBAN forms a homo-dimer and binds to M1-linked Ub chain

ABIN-1 preferentially binds M1- over K48- or K63-linked Ub chains (Fig. 1B). To investigate the structure of ABIN-1 UBAN, and its interactions with Ub chains, we determined crystal structures of ABIN-1 UBAN alone and in complex with M1-linked di-Ub chains (Fig. 2A-C, Table 1). ABIN-1 crystals diffracted to 1.75 Å and contained two coiled-coil homo-dimers in each asymmetric unit of the C2 space group (Fig. 2A, S1A). The two ABIN-1 dimers adopt highly similar conformations (RMSD= 0.87 Å for the superimposition of Cα atoms) and provide two symmetric Ub-binding sites on either side of the dimer (Fig. S1B). Interestingly, mixing ABIN-1 and M1-linked di-Ub in 1:1 and 5:1 ratios prior to crystallization resulted in co-crystals with two different stoichiometries, including 2:2 and 2:1 for ABIN-1: M1-linked di-Ub, that diffracted to 1.95 Å and 3.00 Å resolution, respectively (Fig. 2B & C). Both co-crystals belonged to the *P*2_1_2_1_2_1_ space group and contained one complex consisting of a homodimer of ABIN-1 and one or two M1-linked di-Ub molecules in each asymmetric unit. Despite the difference in stoichiometry, the two complex crystal structures are highly similar in terms of conformation and binding mode (Fig. S1C). In both complex crystal structures distal Ubs bind UBAN through the hydrophobic I44 surface, while the proximal Ub’s interaction with ABIN-1 UBAN is centered on F4 (Fig. 2D). This mode of binding has been previously reported for the UBAN domains of NEMO, OPTN and ABIN-2 with M1-linked Ub chains (Fig. 2E) [21-23].

**Figure 2.**
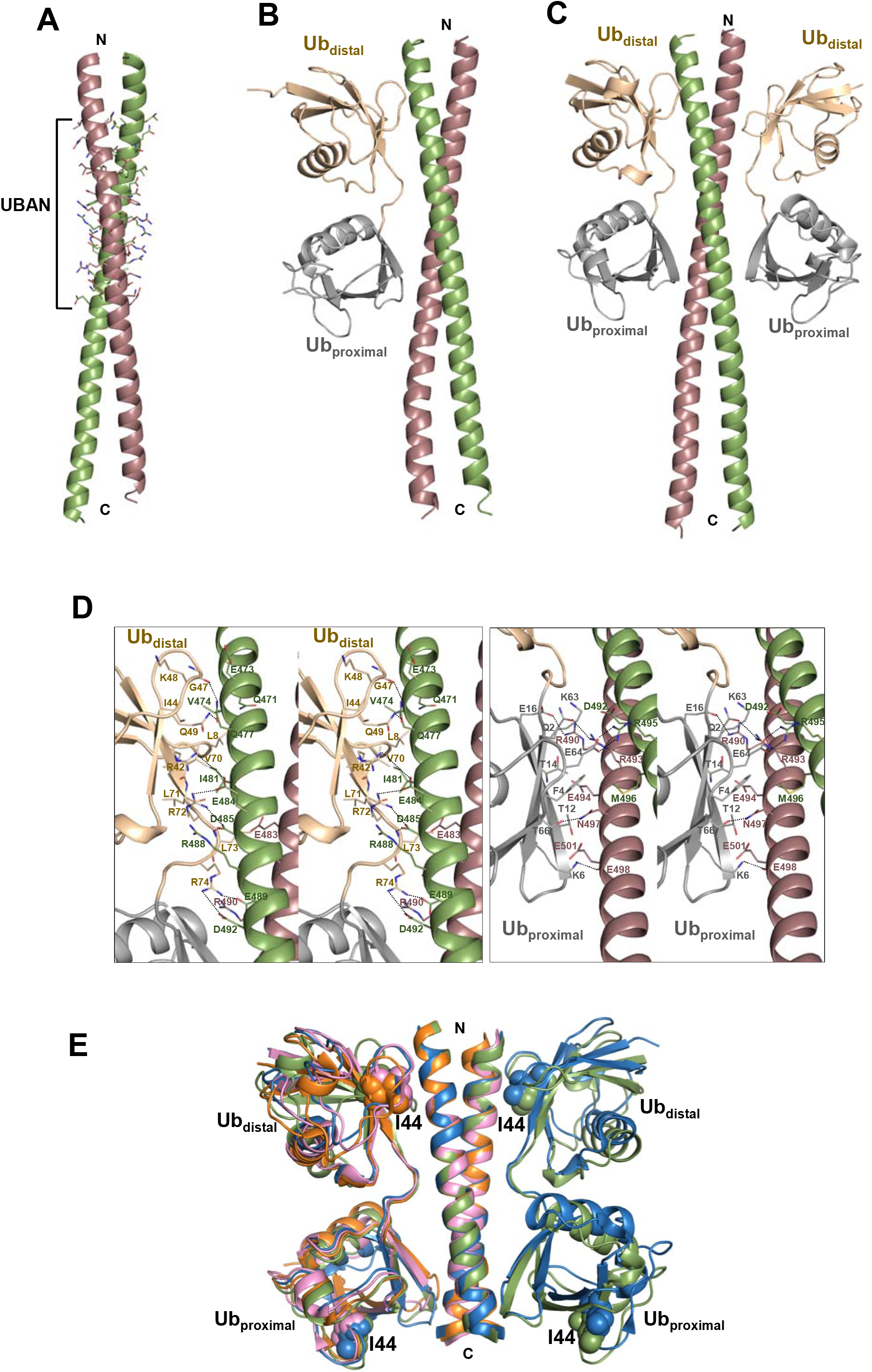
Crystal structures of ABIN-1 (aa 463-532) (**A**) alone, and (**B**), (**C**) in complex with one and two M1-linked di-Ub chains. Residues of UBAN domain are shown as sticks. The two chains of ABIN-1 dimer are colored in green and pink. Distal and proximal Ub moieties in a M1-linked di-Ub chain are colored in light orange and grey, respectively. (**D**) Stereo view of ABIN-1 interactions with distal (left panel) and proximal (right panel) Ub moieties; (**E**) Superimposition of UBAN domains of ABIN-1 (6N6R), ABIN-2 (5H07), NEMO (PDB: 2ZVO) and OPTN (PDB: 3B83) structures in complex with M1-linked Ub chains. The four complex structures are shown in green, orange, blue, and pink, respectively. Ile44 residues in distal and proximal Ub moieties are shown as spheres.

**Table 1.**
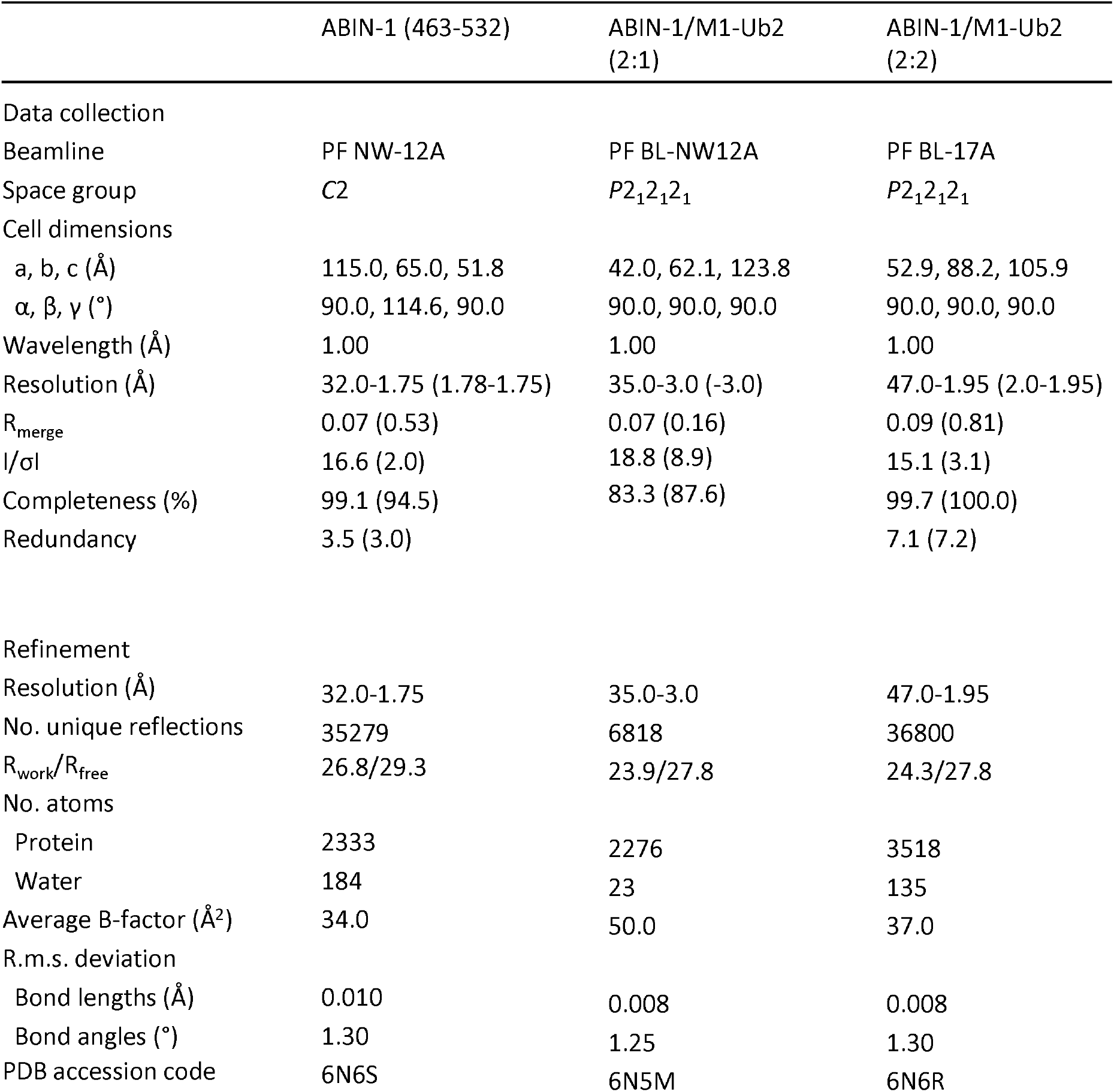
Summary of x-ray diffraction data collection and analyses for ABIN-1, ABIN-1/ M1-Ub2 (2:1 stoichiometry), and ABIN-1/ M1-Ub2 (2:2 stoichiometry) crystals.

### Stoichiometry of UBAN/Ub binding is concentration-dependent

Crystal structures of UBAN domains of NEMO, OPTN and ABIN-2 with M1-linked Ub chains indicate a 2:2 (UBAN: M1-linked di-Ub) stoichiometry. However, insolution methods such as analytical ultracentrifugation and small angle x-ray scattering (SAXS) indicate the existence of an alternative form of NEMO/M1-linked di-Ub complex, in which the two molecules are present in 2:1 ratio [25, 26]. In our co-crystallization of ABIN-1 UBAN/M1-linked di-Ub we obtained both 2:1 and 2:2 stoichiometry by using protein mixtures with 5:1 and 1:1 molar ratios (ABIN-1: M1-linked di-Ub). To further investigate the stoichiometry of complex formation by UBAN and M1-linked di-Ub, we performed isothermal titration calorimetry measurements (Fig. 3A). Interestingly, injection of NEMO or ABIN-1 to M1-linked di-Ub solution resulted in a two-phase binding, starting with a 2:2 stoichiometry. This appears to be due to the abundance of di-Ub molecules in the cell, which readily occupy Ub-binding sites on UBAN domains. In the second phase, stoichiometry of binding changes to 2:1, since further injections provides excess amounts of UBAN. However, in the reverse experiment, where di-Ub was injected to UBAN solution, excess amounts of UBAN in the cell and dynamic interaction of UBAN/M1-linked di-Ub allowed detection of 2:1 stoichiometry, exclusively (Fig. 3B).

**Figure 3.**
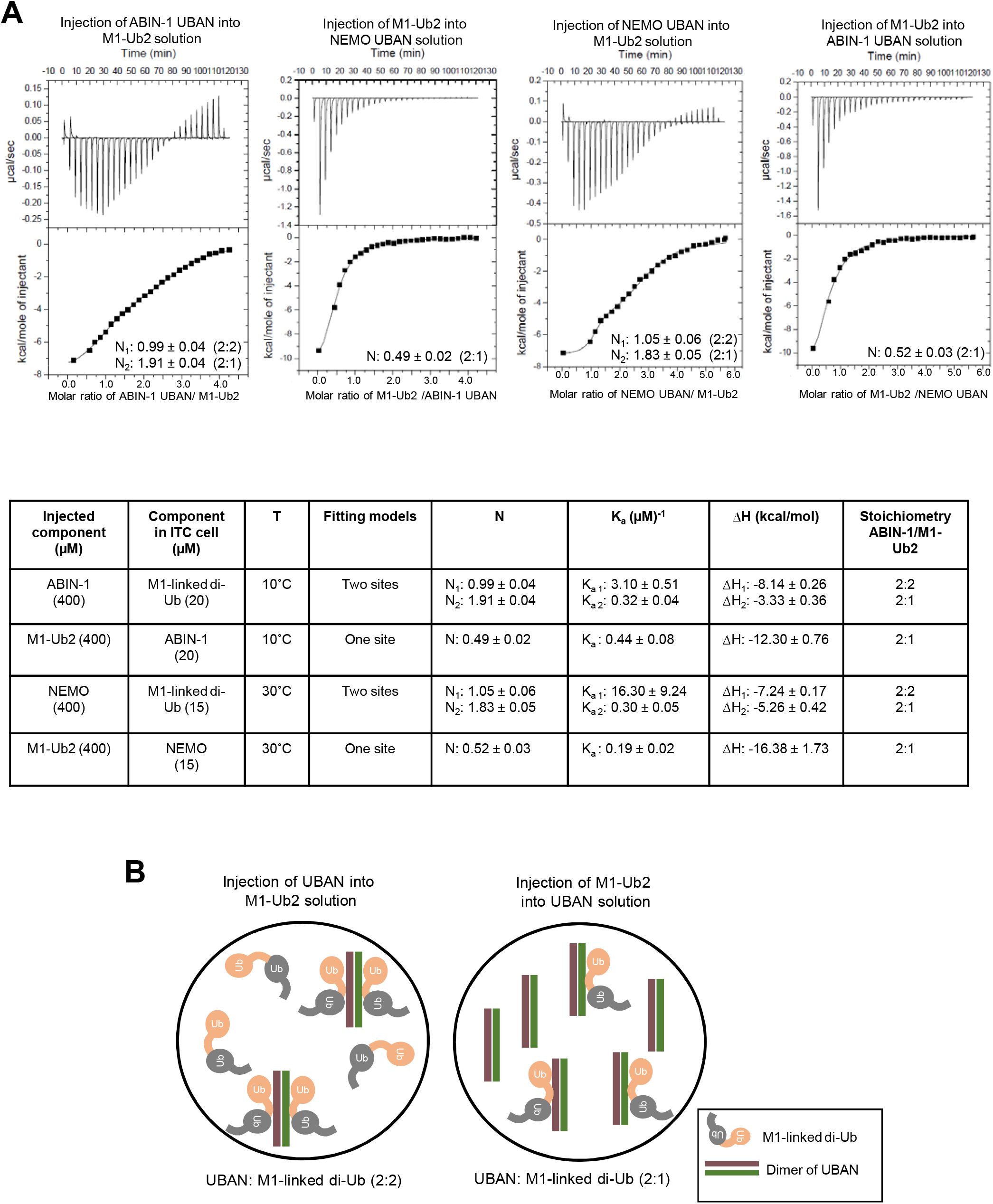
Stoichiometry of UBAN/M1-linked di-Ub binding is concentration-dependent. (**A**) Isothermal titration calorimetry (ITC) measurements and analyses of NEMO and ABIN-1 interactions with M1-linked di-Ub chains; (**B**) schematic representation of UBAN/di-Ub ITC titrations.

### Phosphorylation of UBAN enhances binding to M1-linked Ub chains

To further characterize the Ub-binding properties of UBAN domains, we studied effects of UBAN phosphorylation on UBAN/Ub binding. It has been shown that phosphorylation of S473 in OPTN UBAN enhances its binding affinity towards Ub chains [27]. S473 in OPTN corresponds to the alanine (A303) residue in NEMO and a phospho-mimetic residue (E or D) in ABIN1-3 (Fig. 1A). Since UBAN domains bind Ub in a similar manner, we tested if phosphorylation of these proteins has comparable effects on their binding to M1-linked Ub chains. We measured binding affinities of isolated UBAN domains of OPTN (aa 420-509), ABIN-1 (aa 463-532), and NEMO (aa 250-339) for M1-linked tetra-Ub chains using surface plasmon resonance (SPR) (Fig. 4A, S2). In these measurements, phospho-mimetic OPTN protein with the S473E mutation indicated a K_D_ value of 508 nM, which is ~ 2-fold increase in affinity towards M1-linked tetra-Ub as compared with WT OPTN protein (K_D_ =916 nM). Conversely, ABIN-1 WT is intrinsically phospho-mimetic by containing E484 in the UBAN domain. Therefore, mutation of this residue to alanine (E486A) decreased its binding affinity for M1-linked tetra-Ub (K_D_ of 972 nM and 623 nM for WT and the E486A mutant proteins, respectively). Also, a phospho-mimetic mutation in NEMO UBAN (A303E) increased its binding affinity to M1-linked tetra-Ub chains as indicated by KD values of 562 nM for WT and 362 nM for mutant proteins.

**Figure 4.**
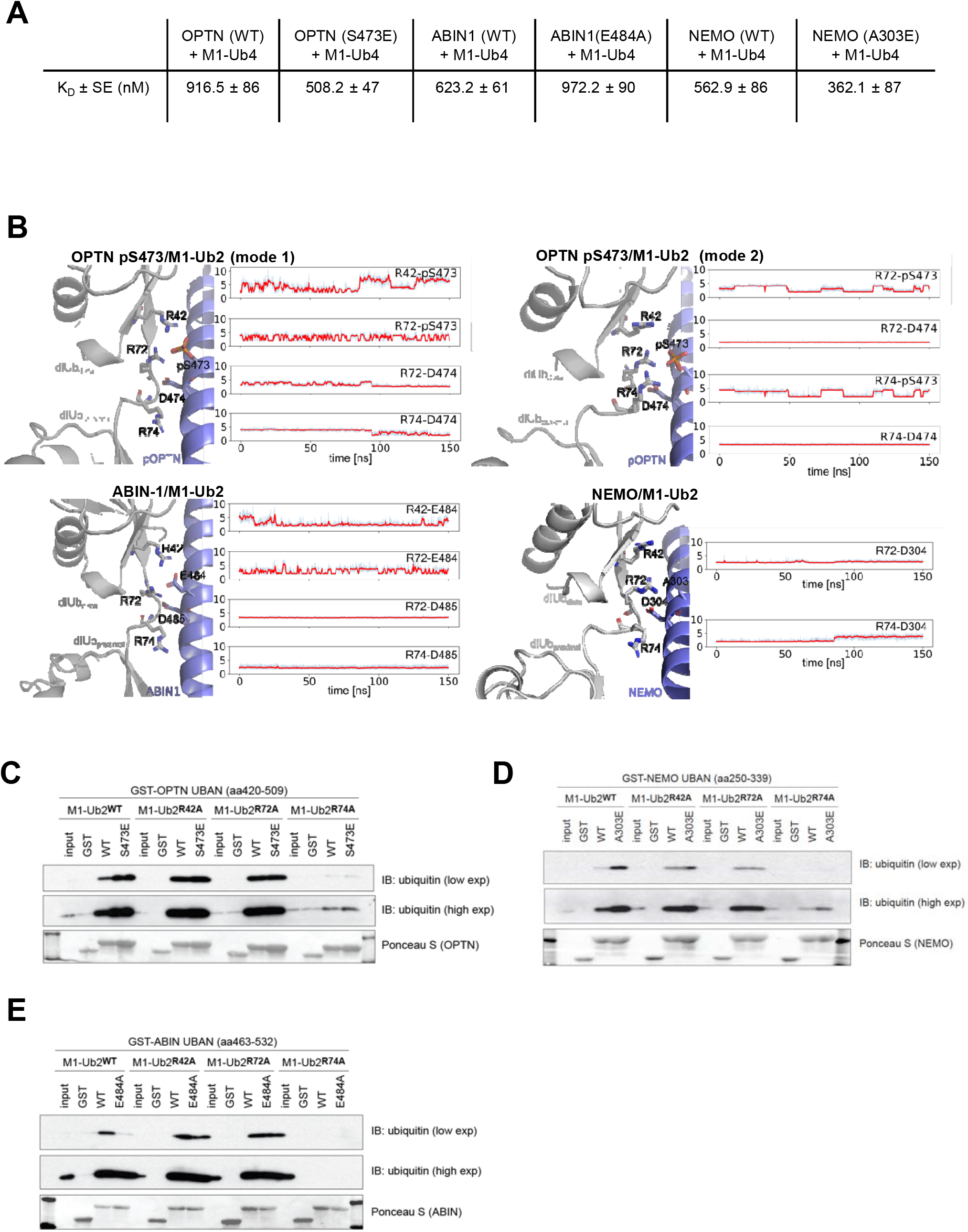
Phosphorylation of S473 increases binding affinity of OPTN UBAN for M1-linked di-Ub chains. (**A**) Binding affinity (K or equilibrium dissociation constants) of UBAN domains of ABIN-1, NEMO, or OPTN (WT or phospho-mimicking) for M1-linked tetra-Ub chains. For each measurement, GST-tagged UBAN was immobilized on the surface of a CM5 chip and ubiquitin chains were loaded over the chip. Each measurement was repeated three times (see also Fig. S2); (**B**) OPTN pS473 interacts with M1-linked di-Ub using distinct, but highly similar modes. If R42 and R72 simultaneously interact with pS473, R72 also interacts with D474, while R74 is not a major partner of UBAN. By contrast, if R72 and R74 interact with pS473, the situation is mirrored and R42 is not a major partner. In ABIN1, the UBAN/M1-linked Ub2 interactions appear highly similar to those of pOPTN, with the phospo-mimetic E484 adopting the role of pS473. In NEMO, in the absence of a charged residue at the phosphorylatable position, the interaction is mainly modulated by D304 and R72. The plots show hydrogen bond distances between hydrogen atoms of arginine (NH_2_) and side-chain oxygen acceptors; Immunoprecipitation of purified UBAN domains of (**C**) OPTN, (**D**) NEMO, and (**E**) ABIN1 proteins with M1-linked di-Ub chains (WT or mutants including R42A, R72A, R74A).

To provide a structural basis for the effect of UBAN phosphorylation on binding to M1-linked di-Ub, we carried out molecular dynamics (MD) simulations [28]. The simulations indicate that R42, R72 and R74 from the distal Ub are the main binding partners of pS473 in the OPTN UBAN (pS473)/M1-linked di-Ub complex (Fig. 4B). We observed slightly different binding modes for the interactions of the two di-Ub molecules with pS473. In one binding mode R42 and R72 interact simultaneously with pS473, although the interaction of R42 with pS473 appears weaker than that of R72 (Fig. 4B, mode 1). In an alternate binding mode, we observed an interaction of R74 guanidino group with the phosphate group (Fig. 4B, mode 2). This mode abrogated any R42-pS473 interaction and required disruption of the highly conserved R74-E478 interface (Fig. S3). Both modes occur in parallel with the strong R72-D474 interaction (Fig. 4B). For NEMO UBAN, our simulations suggest that D304 interacts with M1-linked di-Ub mainly through R72 (Fig. 4B). By contrast, whereas NEMO features a hydrophobic A303 neighboring D304, the corresponding position in ABIN1 is occupied by phospho-mimetic E484. The longer and negatively charged E484 side-chain can interact with R42, and intermittently engages R72. Thus, the interaction pattern is similar to that of pS473 OPTN UBAN (Fig. 3B).

Our *in vitro* pull-down assays further confirm that S473E mutation in OPTN UBAN and A303E mutation in NEMO UBAN increases binding of these proteins to M1-linked di-Ub chains (Fig. 4C, D). This increase in binding is not observed if R42 or R72 of the distal Ub in the M1-linked di-Ub are mutated to alanine. On the other hand, E484A mutation in ABIN-1 decreases its binding to M1-linked di-Ub chains, and R42A and R72A mutations in di-Ub hinder this effect (Fig. 4E). Notably, R74A mutation completely abolishes binding to the UBAN domain of OPTN, NEMO and ABIN-1 proteins.

## Discussion

UBAN-containing proteins play essential roles in various cellular functions, including regulation of NF-κB signaling, protein trafficking, and autophagy, which are mediated by binding to Ub chains through UBAN domains [29]. UBAN domains preferentially bind M1-linked Ub chains, but they also bind K63-linked Ub chains with much lower affinity (Fig. 1B) [21, 24, 30, 31]. Stoichiometry of UBAN/Ub interactions has been a matter of controversy. While most crystal structures of UBAN/Ub complexes demonstrate binding of a UBAN dimer to two Ub chains, in-solution experiments have provided evidence for the formation of a complex with 2:1 stoichiometry (UBAN: diUb) [25, 32]. Our data suggest that although UBAN domains provide two highly symmetrical surfaces on either side of the coiled-coil structure, binding of M1-linked di-Ub chains to one UBAN molecule may not occur, simultaneously, unless significantly high concentration of M1-linked di-Ub chains are available. Thus, the stoichiometry of UBAN/Ub binding is dependent on the abundance of M1-linked di-Ub molecules, which might work as a regulatory mechanism for activation and cellular functions of UBAN-containing proteins.

Mitophagy activates TBK1, which can phosphorylate OPTN at S473 in the UBAN domain [16]. This additional phosphorylation of OPTN boosts its binding affinity to Ub, thereby further driving the mitophagy. By contrast, ABIN-1 natively has a phospho-mimetic (E484) and NEMO has a nonphosphorylatable (A303) residue at the position corresponding to OPTN S473. Correspondingly, ABIN-1 E484A exhibits decreased affinity to diUb, whereas for NEMO A303E affinity increases suggesting OPTN can mimic ABIN-1- or NEMO-like behavior. Our MD simulations also suggest S473 phosphorylated OPTN can adopt ABIN-1-like behavior, engaging R42 in Ub binding. However, while this binding behavior to Ub emphasizes the important role of OPTN pS473 during mitophagy, a similar role for ABIN-1 has not been established.

## Material and Methods

### Construction of plasmids

GST-fusion proteins (UBAN domains of OPTN, ABIN, NEMO and Ub) were cloned into pGEX-4T-1 (GE Life Sciences, Freiburg, Germany) and site-directed mutagenesis was performed by PCR to introduce desired amino acid substitutions.

### Protein expression and purification for pull-down assays

GST fusion proteins were expressed in the *E. coli* strain BL21 (DE3). Bacteria were cultured in LB medium supplemented with 100 μg/mL ampicillin and 0.25 mM ZnSO_4_ at 37 °C in a shaking incubator (150 rpm) until OD at 600 nm reached ~0.5-0.6. Protein expression was induced by the addition of 0.2 mM IPTG and cells were incubated at 16 °C for 16 hours. Bacteria were harvested by centrifugation (4000 rpm) and lysed by sonication in GST lysis buffer (20 mM Tris-HCl, pH 7.5, 10 mM EDTA, pH 8.0, 5 mM EGTA, 150 mM NaCl, 0,1% β-mercaptoethanol, and 1 mM PMSF). Lysates were cleared by centrifugation (10000 rpm) and incubated with Glutathione Sepharose 4B beads (GE Life Sciences, Freiburg, Germany) on a rotating platform at 4°C for 1 hour. After five washes in GST wash buffer (20 mM Tris·HCl, pH 7.5, 10 mM EDTA, pH 8.0, 150 mM NaCl, 0,5% Triton X-100, 0,1% β-mercaptoethanol, and 1 mM PMSF), immobilized proteins were reconstituted in GST storage buffer (20 mM Tris·HCl, pH 7.5, 0.1% NaN_3_ and 0.1% β-mercaptoethanol)[33]. GST was removed from M1-linked di-Ub by cleavage with thrombin. The GST-fused proteins were incubated with thrombin for GST-cleavage in a buffer (20 mM TrisCl, pH 8.5, 150 mM NaCl, 2.5 mM CaCl_2_, 1mM DTT) on a rotating platform at 22°C for 16 hours. Thrombin was inactivated with PMSF.

### GST pull-down assays

Purified proteins (OPTN or ABIN-1) were immobilised on GST beads and combined with 0.5 μg M1-linked di-ubiquitin chains in 500-μL pull down buffer (150 mM NaCl, 50 mM, Tris, pH 7.5, 0.1% Nonidet P-40, supplemented with 5 mM DTT and 0.25 mg/mL BSA)[33]. The proteins were incubated on a rotating platform at 4°C for 16 hours. After five washes with buffer, the proteins were diluted with SDS sample buffer (62.5 mM Tris-HCl pH 6.8, 10% (v/v) glycerol, 2% (w/v) SDS, 0.02% (w/v) bromophenol blue, 5% (v/v) β-mercaptoethanol), resolved by SDS-PAGE and analyzed by immunoblotting with the indicated antibodies.

### Western blotting and Antibodies

For immunoblotting, proteins were resolved by SDS-PAGE and transferred to 0.45-μM nitrocellulose or PVDF membranes. Blocking and primary antibody incubations were carried out in 5% BSA in TBS-T (150 mM NaCl, 20 mM Tris, pH 8.0, 0.1% Tween-20). Secondary antibody incubations were carried out in 5% low-fat milk in TBS-T and washings in TBS-T. Blots were developed using Western Blotting Luminol Reagent (sc-2048; Santa Cruz). The following antibodies were used in this study: anti-ubiquitin P4D1 (#3936; CST) and HRP conjugated goat anti-mouse (sc-2031; Santa Cruz).

### Protein expression and purification for crystallization

ABIN-1 (mouse, aa 463-532) and M1-linked di-Ub were cloned into the pGEX-4T-1 vector (GE-healthcare) and overexpressed as GST-fusion proteins in *E.coli*, BL21 cells. Se-Met containing ABIN-1 protein was expressed in *E. coli* DL41 cells grown in LeMaster medium supplemented with 25mg/L seleno-L-methionine (Wako Pure Chemical). Protein expression was induced by the addition of 0.5 mM IPTG and cells were incubated at 25°C, overnight. Harvested cells were lysed by sonication in PBS buffer and supernatant was applied to a Glutathione Sepharose 4B column (GE-Healthcare). The GST tag was cleaved on the column using thrombin protease and protein was eluted with PBS buffer. Proteins were further purified by gel filtration chromatography using a Superdex 75 column (GE Healthcare) in a buffer containing 20 mM Tris-HCl pH 8.0, and 150 mM NaCl.

### Crystallization, x-ray diffraction data collection, and structure determination

Crystals of ABIN-1 grew in sitting drops containing 20% (w/v) polyethylene glycol 3350 and 0.2 M sodium malonate pH 5.0 in the reservoir solution. Crystals of ABIN-1 in complex with one M1-linked di-Ub were obtained in a condition containing 20% (w/v) polyethylene glycol 3350 and 0.2 M ammonium acetate pH 7.1. Crystals of ABIN-1 in complex with two M1-linked di-Ubs were obtained in a condition containing 30% v/v PEG-MME550, 0.1 M bis-tris pH 6.5, and 0.05 M calcium chloride dihydrate. X-Ray diffraction data were collected at Photon Factory (BL-17A and NW-12A), KEK (Tsukuba, Japan) at 100K, and processed using HKL 2000 [34] or iMosflm. The structure was solved using mono-ubiquitin (PDB: 1UBQ), NEMO (PDB: 3F89) and NEMO/M1-linked di-Ub complex (PDB: 2ZVO) structures as search models for molecular replacement in MOLREP [35]. The model was further built using Coot [36] and refined by the application of amplitude-based twin refinement in REFMAC5 [37, 38]. Data collection and refinement statistics are summarized in Table 1. All structure figures were prepared in PyMOL (PyMOL Molecular Graphics System, Version 1.5.0.5; Schrödinger).

### Surface Plasmon Resonance (SPR)

The SPR experiments were performed using the Biacore S200 instrument (GE-Healthcare). GST-tagged UBAN domains isolated from ABIN-1, NEMO and OPTN were immobilized on the CM5 sensor chip. Various dilutions of M1-linked tetra-Ubs were prepared in the running buffer that contained 10 mM HEPES, pH 7.4 supplemented with 200 mM NaCl, and 0.005% Tween-20. Each experiment was done in triplicate.

### Isothermal Titration Calorimetry (ITC)

ITC experiments were performed using a VP-ITC system (MicroCal) using a buffer containing 20 mM Tris-HCl, pH 8.0 and 150 mM NaCl. The purified NEMO (aa 250-339), ABIN-1 (aa 463-532), and M1-linked di-Ub proteins were degassed in preparation for the experiment. The calorimeter cell and injection syringe were extensively rinsed with buffer. The calorimetric titrations were carried out at 30°C (NEMO) and 10°C (ABIN-1, as the protein was found to be more stable in lower temperature) with a total number of 30 injections spaced 240 seconds apart. For each experiment, ITC data were corrected for the heat of dilution. The Origin software (version 7) was used to analyze the data.

### MD simulations

All systems were parameterized with the Amber ff14SB all-atom force field [39]. For phosphorylated systems, phosphoserines with a net 2^-^ charge were added at the corresponding serine residues of the UBAN domains. Each system was solvated with TIP4 water molecules in a periodic boundary cell with a 10Å solvent buffer, and electrostatically neutralized by adding Na^+^ ions. Specifics for each system are listed in the table below.

| System | Box dimensions<br>(Å) | TIP4 waters | Na<sup>+</sup> | Total atoms |
| --- | --- | --- | --- | --- |
| NEMO:M1 di-Ub | 154.73 92.49 80.76 | 32,501 | 11 | 137,731 |
| pOPTN:M1 di-Ub | 150.10 92.49 80.76 | 31,716 | 21 | 134,389 |
| ABIN1:M1 di-Ub | 86.29 93.28 129.54 | 29,333 | 4 | 124,580 |

In the absence of a 2:2 stoichiometry OPTN:M1 di-Ub structure, we prepared an initial model by threading the OPTN sequence onto the NEMO crystal structure. The systems were minimized with a series of steepest descent and conjugate gradient algorithms by gradually reducing constraints on the protein atoms. Hydrogen atoms were constrained with SHAKE [40]. The time step was set to 1 fs for the initial phase of the NPT (constant N, Pressure, and Temperature) equilibration. After reaching 300 K, the density of the system was equilibrated during a 10 ns NPT (T = 300 K, P = 1.01325 bar) run. The temperature was controlled with the Langevin dynamics method while keeping the pressure constant using the combined Langevin piston Nose-Hoover method. Long-range electrostatic interactions were treated with Particle Mesh Ewald (PME), with a grid spacing of 1 Å. Nonbonded cutoff was set to 12 Å during minimization and heating, and to 10 Å during the NPT equilibration and production simulation. The integration time step for the final stages was increased to 2 fs. All simulations were performed with openMM on Kepler K20 and GTX 1080 GPUs [41].

### Accession numbers

Atomic coordinates and structure factors of the ABIN-1 alone, and in complex with one and two M1-linkd di-Ub chains are deposited in the PDB with accession codes PDB: 6N6S, 6N5M, and 6N6R, respectively.

## Acknowledgments

We would like to thank Dr. Masato Kawasaki and staff of the Photon Factory BL-17A and AR-NW12A beamlines for their support with the X-ray diffraction data collection. This work was supported by grants from Japan Society for Promotion of Science (JSPS), DFG (SFB 1177 on selective autophagy), the Cluster of Excellence “Macromolecular Complexes” of the Goethe University Frankfurt (EXC 115), LOEWE grant Ub-Net and LOEWE Centrum for Gene and Cell Therapy Frankfurt. L.H. is supported by the European Molecular Biology Organization (EMBO) long-term postdoctoral fellowship.

---

**Figure S1.**
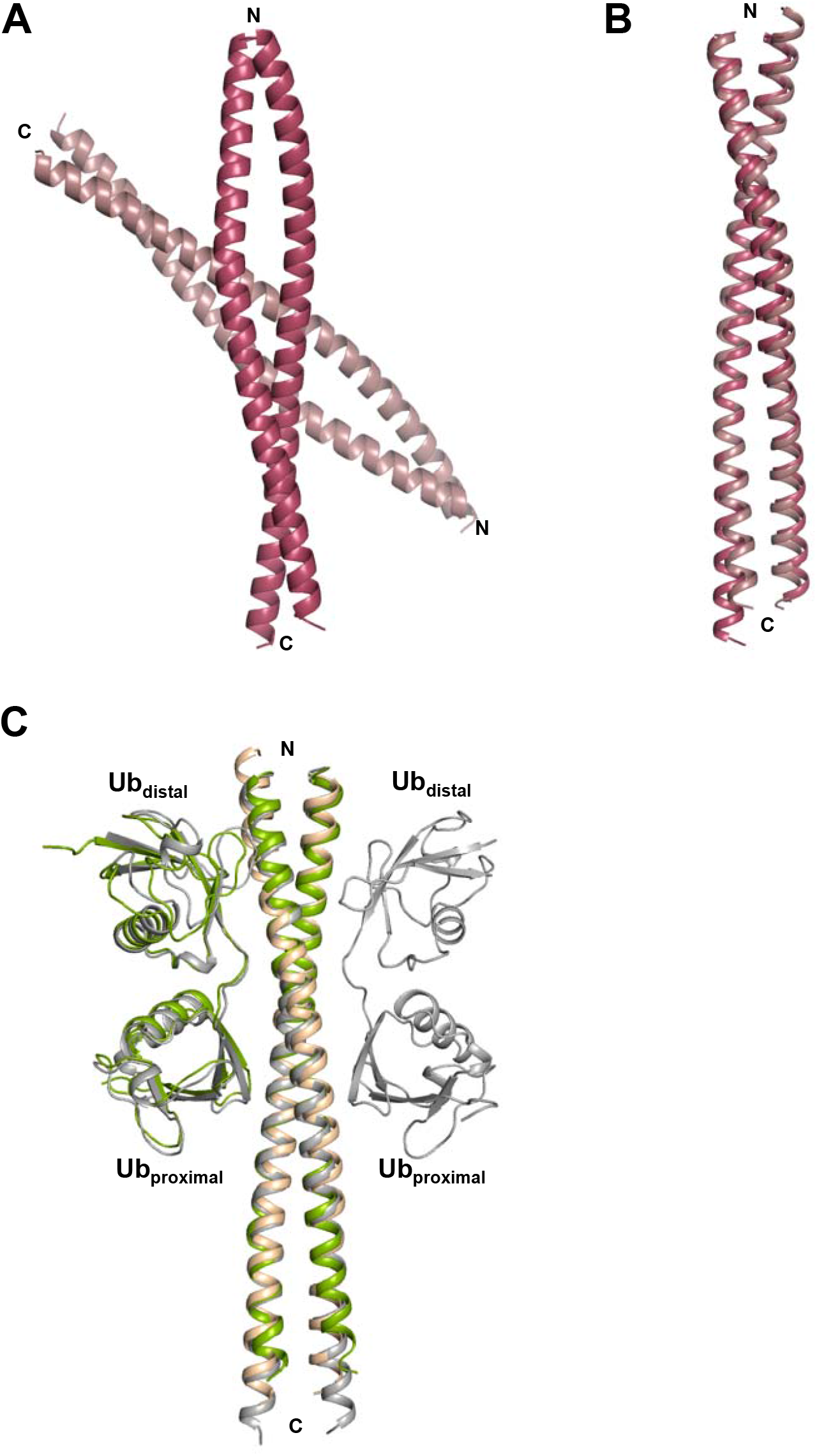
Crystal structures of ABIN-1 alone and in complex with M1-linked di-Ub chains. (**A**) There are two ABIN-1 dimers in the asymmetric unit of the C2 crystal; (**B**) the two ABIN-1 dimers are well superimposed as indicated by RMSD of 0.87 Å for the superimposition of Cα atoms (residues 465 to 530); (**C**) superimposition of ABIN-1 structure in the free form (light orange) and in complex with one (green) or two (gray) M1-linked di-Ub molecules.

**Figure S2.**
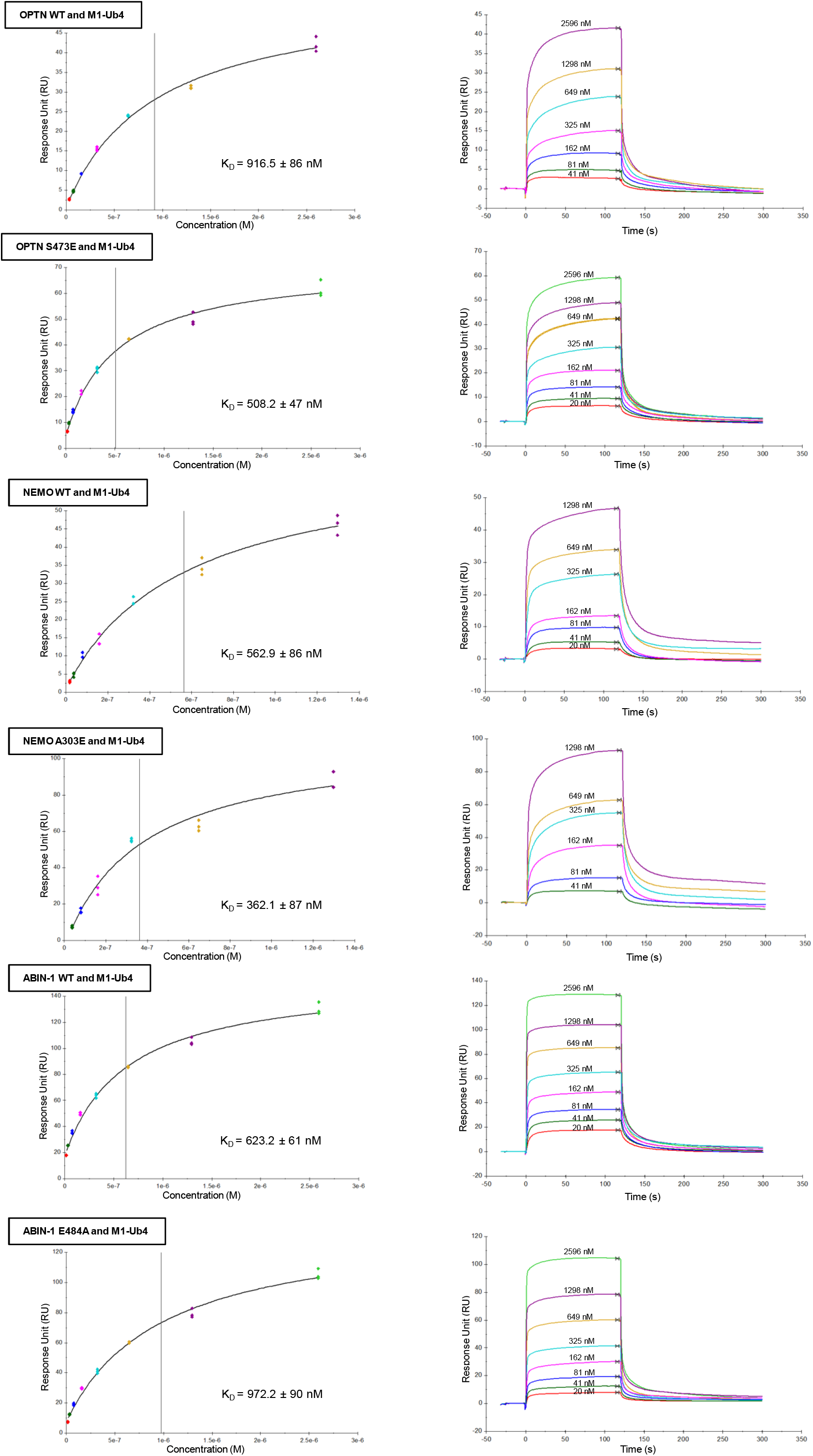
Surface plasmon resonance (SPR) (left) Equilibrium fitting and (right) sensograms for wild-type (WT) and mutated UBAN domains of OPTN, NEMO and ABIN-1 proteins. Each measurement was don in triplicate.

**Figure S3.**
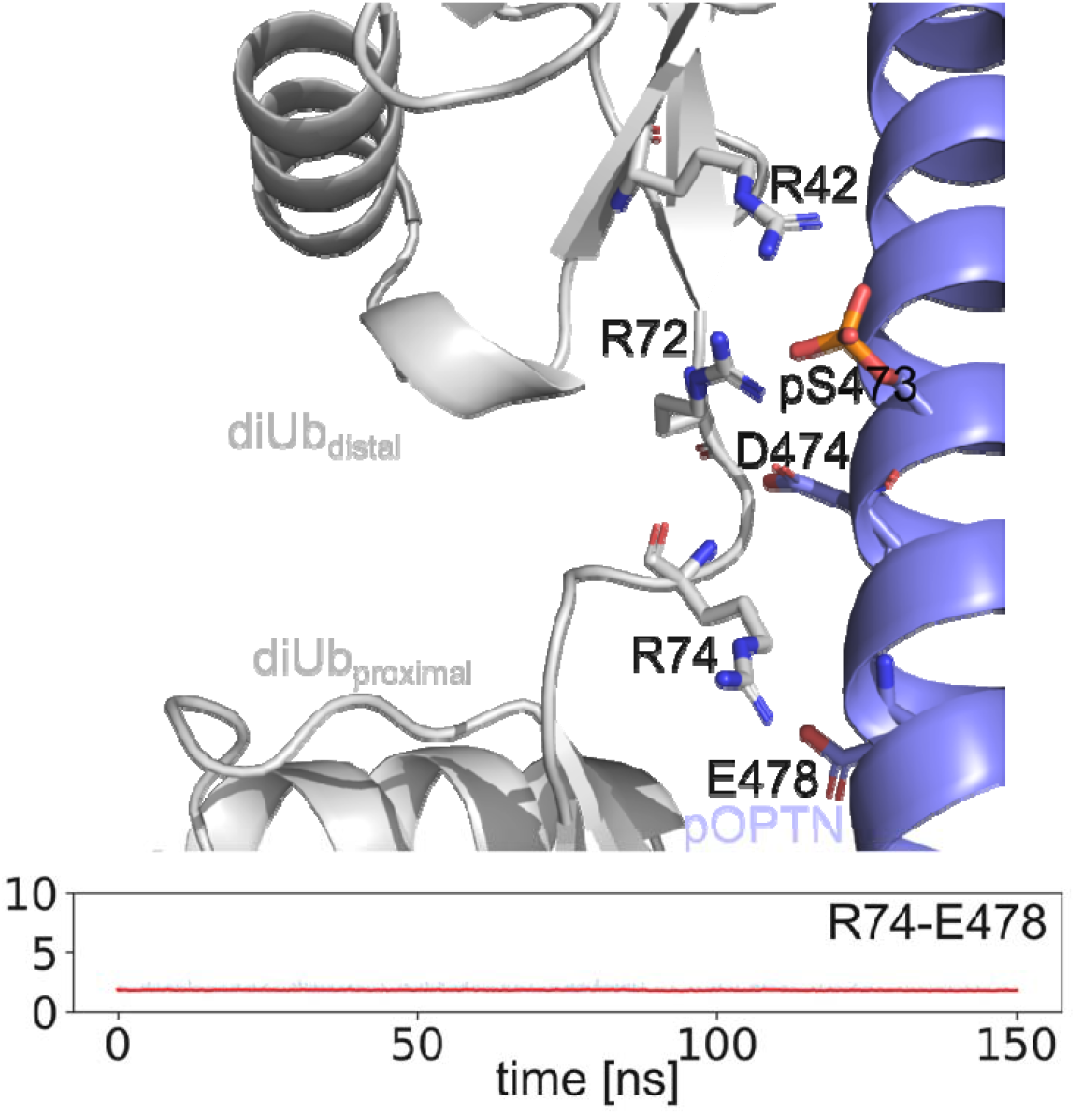
Interaction of the R74 guanidino group with the phosphate group in the OPTN UBAN (pS473)/M1-linked di-Ub complex (Fig. 4B, mode 2) disrupted interaction of R74 (NH_2_) hydrogen atoms with the side-chain carboxyl oxygen of E478. The R74-E478 is highly conserved in UBAN.

